# The Dark Ecology Dataset: Measurements of Biological Activity in US Weather Radar from 1995 to 2025

**DOI:** 10.64898/2026.06.20.733536

**Authors:** Daniel Sheldon, Kevin Winner, Iman Deznabi, Garrett Bernstein, Pankaj Bhambhani, Tsung-Yu Lin, Peter Desmet, Adriaan M. Dokter, Kyle G. Horton, Cecilia Nilsson, Benjamin M. Van Doren, Andrew Farnsworth, Frank A. La Sorte, Subhransu Maji

## Abstract

The US NEXRAD radar network has monitored the aerosphere over the US and its territories continuously since the 1990s and archived nearly 300 million radar volume scans. These data contain a wealth of information about the movements of birds, bats, and insects. Historically, this biological information was difficult to access due to the amount of data and challenges in analyzing it. In the last 15 years, fueled by computational and methodological advances, large-scale aeroecology research has blossomed. However, comprehensive analyses of the NEXRAD archive remain very costly. We collected measurements from every volume scan in the NEXRAD archive—nearly 300 million data files total—to assemble a dataset of aerial biological activity over the US from 1995 to 2025. The core data are vertical profiles, which summarize biological activity at different heights above the radar station for each volume scan. We also provide time series data products that aggregate vertical profiles to point measurements at radar stations across time. These data products can support a range of aeroecology analyses at significantly reduced effort.

## Background & Summary

It was first discovered during World War II that early military radars could detect birds, thus offering an unexpected tool for ornithological research [1, 2]. In the decades that followed, radar technology has developed and advanced significantly, with broad applications in defense, aviation, and meteorology, as well as in biological sciences, becoming a valuable tool for studying birds [3–5], insects [6–8], and bats [9]. Today, radars are essential tools in the emerging discipline of aeroecology, the study of organisms in the aerosphere [10]. The deployment and maintenance of large networks of weather surveillance radars has been particularly impactful. These networks provide continuous monitoring of the atmosphere over large spatial scales and long time periods, making it possible to measure patterns and movements of flying organisms across unprecedented spatial and temporal extents [11–59].

The US NEXRAD (Next-Generation Radar) network [60], also known by the technical name WSR-88D (Weather Surveillance Radar, 1988, Doppler), is particularly noteworthy due to its size, uniformity, and standardized data collection. The network of 158 radar stations in the US and its territories or military installations abroad is one of the largest in the world and has archived data in a consistent format since its installation in the 1990s [61], which makes it one of the largest existing archives of animal movement data. NEXRAD data have transformed how we study bird migration at the largest scales, including nocturnal migration [13, 14, 21–23, 29, 30, 32, 34, 43, 46, 62, 63], stopover behavior [15, 18, 28, 41, 49, 51, 64], waterfowl distributions [65], communal roosts [66–78], demography [27], impacts of artificial light [25, 26, 36, 37, 44, 50], phenology [17,20,39], population declines [38], and migration forecasts [31,42,79], as well as similar patterns for bats [58, 59] and insects [52–55].

Methodological and computational advances over the last 15 years have significantly propelled aeroecology research using NEXRAD data. Historically, access to biological information in NEXRAD data was difficult due to the volume of data, challenges in accessing the radar data themselves, and the lack of algorithms to discriminate and measure biological quantities of interest. Many studies were limited in scope due to computational demands or the need to manually view and interpret images. Advances that have helped overcome these barriers include new conceptual frameworks and algorithms for extracting biological information [15,16,80–84], AI methods to detect and classify biological phenomena [85–88], the development of user-friendly open-source software tool kits [89–91], and community efforts to create and grow the field of aeroecology [10, 19, 92–96]. Another major development was the move in 2016 to host the NEXRAD archive on the Amazon Web Services (AWS) cloud computing platform [61]. It is difficult to overstate the importance of this change, which allowed researchers immediate, on-demand access to archived NEXRAD files and opened the possibility and power of highly scalable cloud computing workflows to analyze the data.

Despite these advances, unlocking the full potential of the NEXRAD archive remains a costly endeavor. Each radar station systematically scans the surrounding volume of atmosphere every 4–10 minutes and archives the results as a volume scan data file or “volume scan”. The archive contains 298 million volume scans comprising 0.84 PB (petabytes) of data. Our workflow, representative of a typical biological analysis, takes an average of 11 seconds per volume scan, which would take 104 years of compute time on a single CPU. These computations are too costly and difficult for most individual research groups. Since many aeroecology analyses rely on similar measurements extracted from the radar data, there is considerable motivation to create and publish datasets of biological activity extracted from NEXRAD data that can be used by a broader community of users, to reduce redundancy and subsequent waste and also to promote consistency and standards within the community.

We present a dataset of animal occurrence and aggregate movement, “biological activity”, in the air, extracted from the complete NEXRAD archive from 1995–2025. Our core data product consists of one vertical profile of biological activity for each radar volume scan in the archive. Vertical profiles summarize the density and velocity of objects, such as animals, that scatter radio waves (“scatterers”) in different elevation bands above the radar station (see Figure 1). They have long been used in both meteorology and aeroecology, especially to measure broad-scale phenomena like nocturnal bird migration and insect movements [89]. We use MistNet [86], a deep neural network developed by our research group, to discriminate biological scatterers from precipitation in historical data, and apply additional filtering steps to remove unwanted scatterers such as ground clutter.

**Figure 1:**
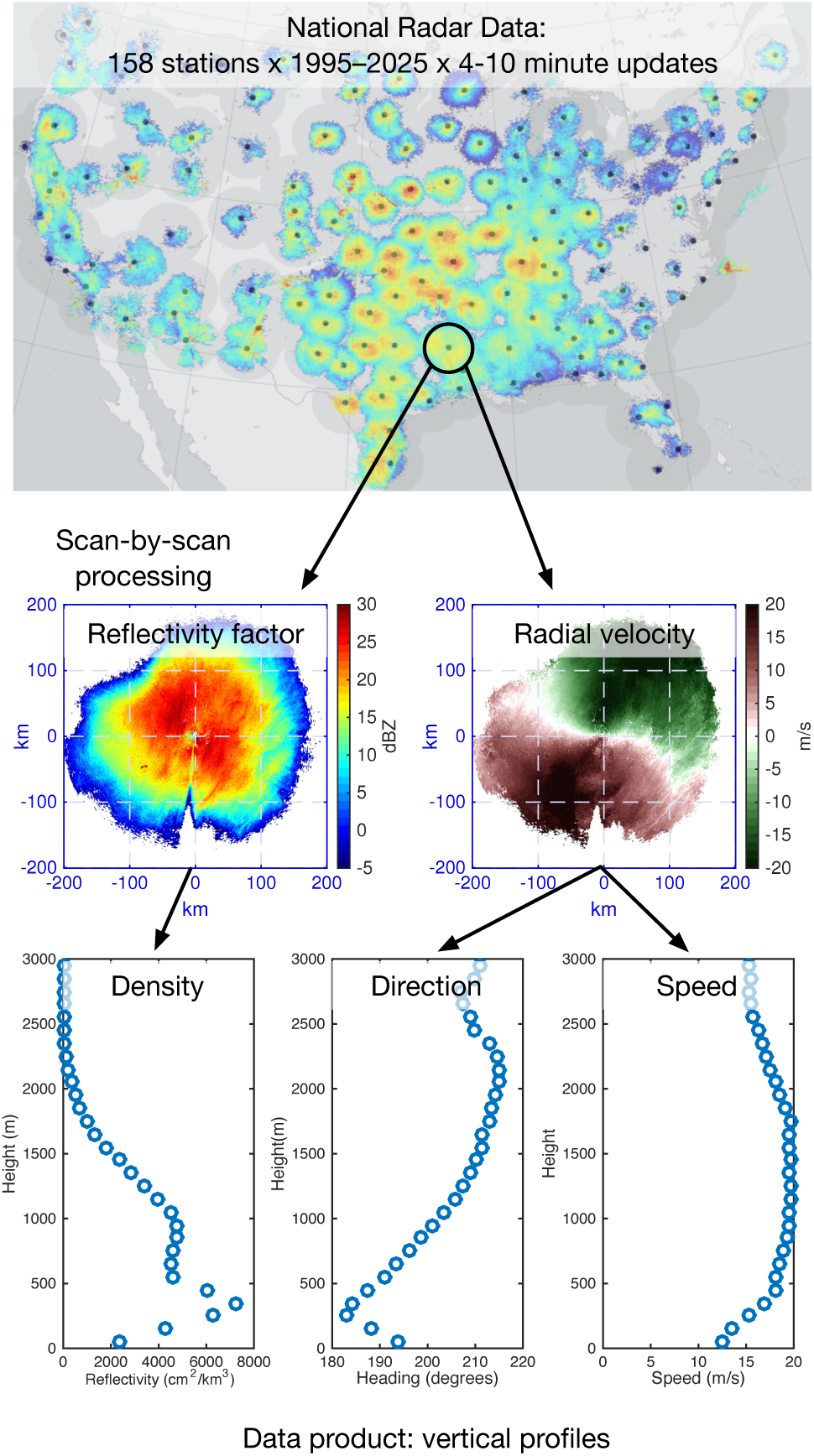
Overview of data preparation workflow for vertical profiles. Each radar volume scan in the archive was processed to create a vertical profile summarizing biological activity at different heights above the radar station.

Vertical profiles are important because they compactly summarize biological activity: our full dataset is about 250 GB, more than 3000 times smaller than the source data, but preserves biologically significant variation across elevations. Vertical profiles can also be integrated vertically to obtain point measurements summarizing the biological activity above different radar stations. We provide several derived data products consisting of time series of point measurements. Our most compact data product is a daily time series of point measurements for all radar stations, with one data file per year and a total size of only 160 MB. These data products can fuel a range of aeroecology analyses at significantly reduced effort. Our collection of data products is named the *Dark Ecology Dataset*, in reference to the use of radar to study the largely unseen world of aerial organisms.

A related effort recently published biological vertical profile data from 141 European weather radars with varying temporal coverage [97]. This dataset is similar in spatial scale to Dark Ecology data but covers a shorter total period (2008–2023) and has more variability due to the relative heterogeneity of radar hardware and practices across Europe compared to those in the US. In the future, standardized datasets of biological activity from weather radar networks in different regions of the world could facilitate globalscale aeroecology research.

## Methods

Our main workflow goal was to process each radar volume scan in the archive to construct a vertical profile (see Figure 1). We built on methods to construct vertical profiles from our prior research studies [21–23, 29, 31, 32, 34, 39, 41–44, 81, 86] and implemented them in the WSRLIB MATLAB toolbox [91]. At a high level, the methods first apply filtering steps to remove unwanted scattering volumes in each volume scan and then aggregate measurements to collect summaries of density and velocity for each height bin. Our method produces very similar results to methods developed and used in a related line of work [16, 27, 30, 33, 35, 38, 45–47] and implemented in the bioRad R package [89]. We provide a detailed comparison between the two approaches in the technical validation section.

To manage the large amount of computing resources to process nearly 300 million data files, we developed cloud computing workflows to construct vertical profiles for many volume scans in parallel. After constructing the vertical profiles, we created aggregated data products by vertically integrating profiles to obtain point measurements and then assembling point measurements into time series. Below, we provide details of the source data, our method for constructing vertical profiles, the cloud computing workflows, and the aggregated data products.

### Source Data

We obtained Level II NEXRAD data files from the unidata-nexrad-level2 S3 bucket on Amazon Web Services [98], where it is freely accessible as part of NOAA’s Open Data Dissemination program (formerly the Big Data Initiative) [61]. There is one data file per radar volume scan (or “scan”), during which a single radar station conducts a three-dimensional scan of the surrounding atmosphere. Volume scans are described further below and illustrated in Figure 3. Each data file contains gridded data products in a compressed binary format with file name indicating the radar station and time of collection. There were a total of 176 WSR-88D stations with data available at some point during the period 1991–2025. Of these, 158 stations were operational at the time of writing, including 143 in the contiguous US, 7 in Alaska, 4 in Hawaii, 1 in Puerto Rico, 2 in South Korea, and 1 in Guam. See Figure 2 and Supplementary Table S1. We excluded data from 15 transient or test-bed stations (call signs DAN1, DOP1, FOP1, KABQ, KCRI, KDOG, KILM, KOUN, KRMX, KUNR, LPLA, NOP3, NOP4, ROP3, ROP4) with fewer than 3 years of data or 100,000 scans. We included data from three stations that are no longer operational: KJAN (Jackson, MS) operated from 1995–2004 before being replaced by KDGX (Brandon, MS), which is 10 km from the original KJAN location [99]; KLIX (New Orleans, LA) operated from 1995–2003 before being moved 59 km to Hammond, LA in 2024 and renamed KHDC [100]; RODN (Okinawa, Japan) archived data from 1996–1999 and 2017–2023 before being retired [101]. We began our analysis with the year 1995, when most stations came online. Our final dataset therefore includes data from 1995–2025 for 161 stations, including the 158 currently operational stations together with KJAN, KLIX, and RODN, for a total of 298.3 million scans and 841 TB of storage. The average number of scans per station is 1.9 million over an average period of 29.2 years, representing a scan every 8.1 minutes on average—we note that this number overestimates the scanning time during continuous operation because stations were not always operational.

**Figure 2:**
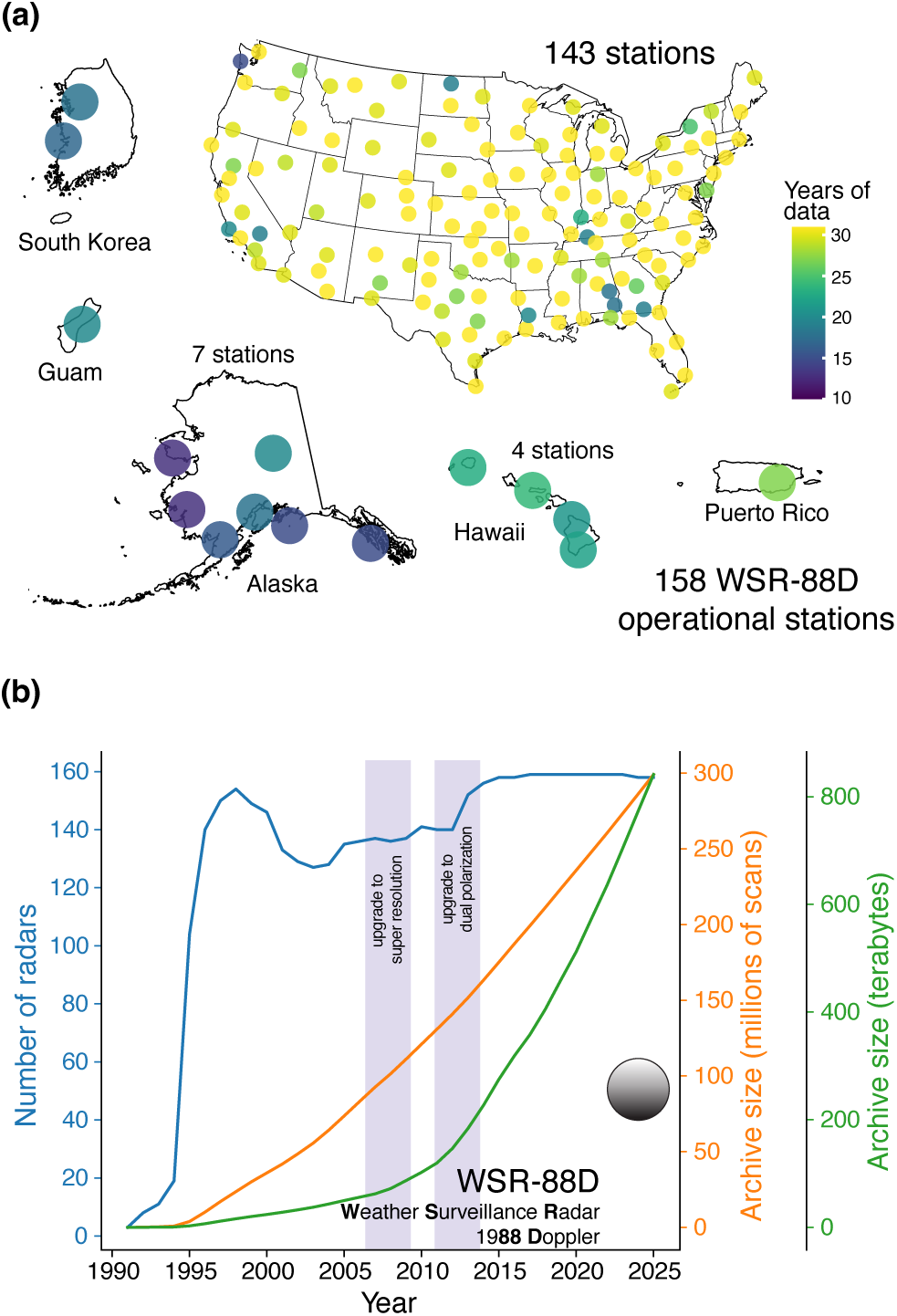
NEXRAD radar locations and archive size. Panel **(a)** shows the locations of the 158 currently operational radar stations in the US and its territories. Panel **(b)** shows the number of active stations and cumulative archive size by year.

**Figure 3:**
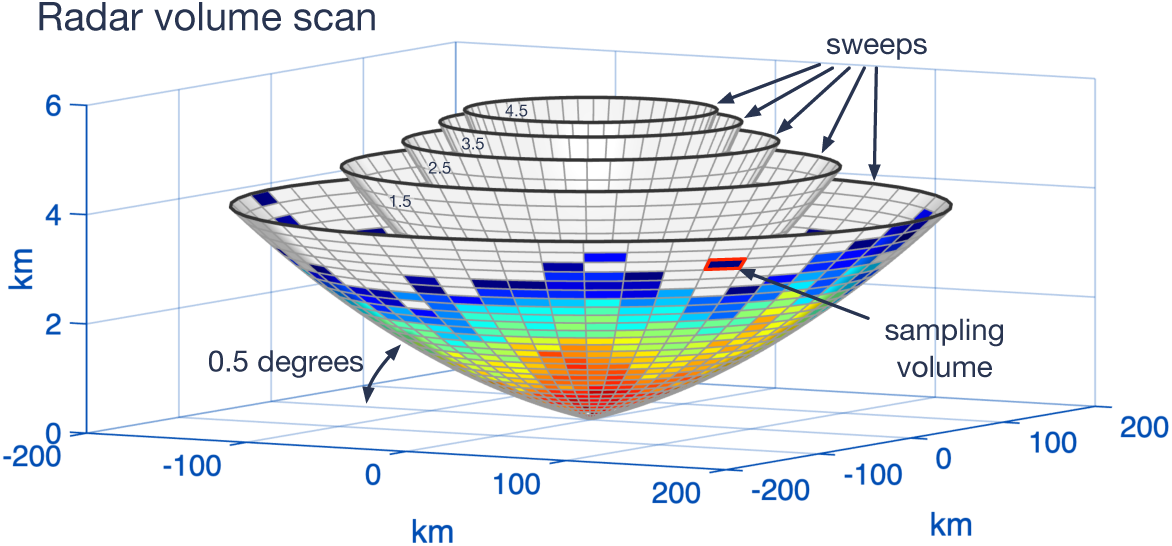
Illustration of radar volume scan. The radar conducts a sequence of sweeps, each time rotating the antenna 360*^◦^*at a fixed elevation angle to collect data along an approximately cone-shaped surface, which is stored in a two-dimensional polar grid. In clear air operations, sweeps are commonly near 0.5*^◦^*, 1.5*^◦^*, 2.5*^◦^*, 3.5*^◦^* and 4.5*^◦^*, as shown here. This illustration truncates the range of each sweep and reduces its resolution for clarity. Sweeps are curved slightly upward relative to the ground, shown here as flat, due to Earth’s curvature—the beam itself typically bends downward slightly due to refraction, but not as quickly as the ground curves away from the beam.

During a volume scan, the radar performs a sequence of sweeps to collect gridded data in three-dimensional polar coordinates. The geometry of a volume scan and its components, sweeps and sampling volumes, is illustrated in Figure 3. During one sweep, the antenna rotates about the vertical axis at a fixed elevation angle to measure scattering of radio waves along an approximately cone-shaped surface of the atmosphere extending to a maximum range of 460 km, but sometimes shorter depending on the data product and elevation angle. To sample the full volume, the radar conducts sweeps at increasing elevation angles, typically ranging from 0.5*^◦^* to as high as 19.5*^◦^*, depending on the operating mode. Three to six data products are collected per sweep and stored as two-dimensional arrays with entries corresponding to sampling volumes at a given range and azimuth relative to the antenna and at the elevation angle determined by the sweep. Three “legacy” data products—reflectivity factor (*Z*), radial velocity (*v_r_*) and spectrum width (*σ_w_*)—are available for the entire archive. Reflectivity factor is related to the density of scatterers in the sampling volume, while radial velocity measures the aggregate velocity of scatterers in the direction towards or away from the radar, and spectrum width is a measure of the variability of radial velocity within the sampling volume. Figure 1B shows reflectivity factor and radial velocity for a single sweep projected into 2D Cartesian coordinates. In 2012 and 2013, NEXRAD sites were upgraded to dual polarization technology and three additional “dual-pol” products became available. Our goal was to measure biological activity in a consistent way across the full duration of the archive, so we did not use dual-pol data for any steps of our processing or filtering. However, our MistNet algorithm used to filter precipitation was trained with the assistance of dual-pol data; see the Meteorological Scattering paragraph in Reflectivity Filtering section below.

When extracting biological measurements, we converted reflectivity factor, as provided in the source data, to *reflectivity* [80], which is an interpretable unit for biological applications. Conceptually, reflectivity factor is obtained from reflectivity by assuming scatterers are small spherical water drops, which is useful for meteorological applications but not biological ones. Reflectivity (*η*, cm^2^ km*^−^*^3^) measures the total scattering area, or summed radar cross-section (RCS, cm^2^) of all scatterers, per unit volume, where RCS measures how reflective an object is to radio waves. For biological scatterers, reflectivity is directly related to the number and physical properties of the organisms. It can be converted to count units by dividing by the estimated RCS for a particular type of animal [80,102] (for example, 11 cm^2^ is frequently used as a representative value for birds [16, 31]) and further to biomass if the average mass of an individual can also be supplied. Because these conversions depend on the types of animals present and their properties, which may vary across our dataset and be difficult to estimate precisely in practice, we left our measurements in units of scattering area; they may be further converted as needed for specific applications.

### Vertical Profiles

We processed the Level II volume scan files to create vertical profiles using the WSRLIB MATLAB toolbox [91], which was developed by our research group and uses NASA’s Radar Software Library (RSL) [103, 104] to read binary Level II files. Code to recreate the vertical profile for any volume scan in the archive is available using the cajun function in WSRLIB Version 1.0.0.

The source data and vertical profiles are illustrated in Figure 1. Vertical profiles are constructed by grouping sampling volumes into 100 m bins based on height above the radar station and then computing aggregate measurements of reflectivity and velocity for each height bin. We included sampling volumes from five sweeps with elevation angles closest to 0.5*^◦^*, 1.5*^◦^*, 2.5*^◦^*, 3.5*^◦^* and 4.5*^◦^* and within 50 km ground distance from the radar station and up to a height of up to 3000 m above the radar, below which most biological activity occurs [16,105–107]. The maximum range of 50 km was chosen to ensure good coverage up to 3000 m using elevation angles up to about 4.5*^◦^*. Sampling volume coordinates were based on point at the center of the range bin for each azimuth and elevation angle.

The procedures to construct profiles for velocity and reflectivity are slightly different, with the following high-level processing steps:

1. Compute velocity profiles following the Bayesian velocity-profiling algorithm of Sheldon et al. (2013) [81].
2. Apply several filtering steps to exclude signals from unwanted scatterers such as ground clutter or precipitation, or from sampling volumes that are not adequately measured due to beam blockage.
3. Compute reflectivity profiles by averaging the reflectivity of non-excluded sampling volumes in each height bin.

In addition to the main reflectivity and velocity measurements, several derived or diagnostic variables are also computed for each height bin—these are described in detail in the data record descriptions. We now discuss each of the high-level processing steps in more detail.

### Velocity Profiles

Radial velocity is the component of scatterer velocity that is directed towards or away from the radar antenna. Velocity orthogonal to this direction is unmeasured. However, by making assumptions about the movement of scatterers, it is possible to reconstruct full velocity vectors. Such methods are common in meteorology and known as *velocity-azimuth display* methods or *velocity volume profiling* (VVP) [108, 109]. The *uniform velocity model*, which assumes velocity is constant across space at a specified height above the ground, is appropriate for both meteorological and biological scatterers, for which velocities tend to vary significantly with elevation but be fairly uniform within the radar domain at the same elevation. By measuring the radial velocity component at different azimuths, these methods can recover the true underlying velocity vector (assumed uniform) in each height bin. The uniform velocity is widely used in meteorology and aeroecology and has been observed to outperform velocity models with more parameters [110]. In the case of heterogeneous velocities, e.g., from birds flying in different directions, or from a mixture of scatterer types, the retrieved velocity measurement represents an aggregate across the height bin.

We used the Bayesian velocity profiling algorithm of Sheldon et al. (2013) [81] as implemented in WSRLIB by the epvvp function. This algorithm handles the issue of velocity *aliasing* together with velocity profiling. Aliasing is a fundamental ambiguity in Doppler velocity measurements, where velocities with magnitude above the Nyquist limit are folded back into the observable range. The epvvp algorithm handles aliasing in a Bayesian context through the use of a wrapped normal likelihood function, which models the physical process of aliasing. The algorithm also uses a prior distribution to encourage smoothness across height bins, which facilitates inference with limited data by sharing information across height bins. Estimation is performed using the expectation propagation algorithm [111] for approximate Bayesian inference.

The primary outputs of velocity profiling are the east-west velocity component (*u*) and north-south velocity component (*v*) inferred for each 100 m height bin. We also computed the direction and speed of movement as derived directly from *u* and *v*, and reported the root-mean squared error (RMSE) of the VVP fit to the uniform velocity model within each height bin. See the data record descriptions for more details.

### Reflectivity Filtering

Before constructing reflectivity profiles, we applied filtering steps to remove radar clutter, sampling volumes affected by radar beam occultation, and precipitation. In this section we describe how we identified these phenomena. Details of applying these filters are described below.

### Static Clutter Maps

We built static clutter maps for each radar station to determine sampling volumes that are consistently affected by scattering from unwanted phenomena, especially ground-based objects such as trees, buildings, or mountains. For each station and year from 1995 to 2020, we computed the probability of detection (POD) for 30 clear air scans from the month of January, representing a baseline period that is expected to be mostly free of biological scattering. We processed the first 300 January scans starting Jan 2 00:00 UTC to avoid New Year’s Eve disturbances [112]. For each scan, we computed reflectivity factor on a fixed polar grid of 1*^◦^* by 500 m to a range of 150 km at elevation angles 0.5*^◦^*, 1.5*^◦^*, 2.5*^◦^*, 3.5*^◦^* and 4.5*^◦^* with values clipped to at most 35 dBZ. We populated measurements for each cell in the fixed polar grid using the nearest actual measurement, i.e., using nearest neighbor interpolation. We then selected the 30 scans with lowest total reflectivity factor as representing clear conditions. From these scans, we computed POD@10dBZ (“probability of detection at 10 dBZ”) for each grid cell, which is the fraction of scans for which the reflectivity factor exceeded 10 dBZ. A grid cell was marked as clutter if POD@10dBZ was 30% or more in two or more years (to help mitigate false positives due to actual atmospheric phenomena in a single year).

### Occultation Maps

Occultation occurs when sampling volumes are “shadowed” by terrain features or other objects closer to the radar. We built static occultation maps for each radar station using the method of Krajewski et al. (2006) [113], which uses a digital elevation map to determine sampling volumes that are completely or partially blocked by terrain features. We calculated the percentage of occultation for each sampling volume on a fixed polar grid of 1*^◦^* by 1 km out to 150 km and for increasing elevation angles starting at 0.5*^◦^* in increments of 0.5*^◦^*until no occultation was present. We then created a binary mask, marking a sampling volume as blocked if the occultation was at least 10 %.

### Meteorological Scattering

A long-standing challenge for biological analysis of weather radar is the identification and removal of precipitation. To identify precipitation we used MistNet [86], a deep neural network for weather radar data. As input it takes a radar volume scan rendered as a multidimensional array in three-dimensional Cartesian coordinates, including the legacy products of reflectivity factor, radial velocity, and spectrum width for the five elevation angles closest to 0.5*^◦^*, 1.5*^◦^*, 2.5*^◦^*, 3.5*^◦^* and 4.5*^◦^*. MistNet outputs a mask predicting which cells in the three-dimensional grid are precipitation vs. biology. Because it uses only legacy products to make its predictions, MistNet can be applied to all historical NEXRAD volume scans.

MistNet was trained with noisy labels derived from dual-pol data products and validated using human annotations of precipitation vs. biological scattering. The training and validation data were focused on time periods containing bird migration but were extensive enough to include many cases of bats and insects. Noisy labels were obtained by thresholding the copolar correlation coefficient (*ρ*_HV_) at 0.95, which is effective at discriminating precipitation (values close to 1) from biological scatterers [27, 83, 114]. The training data included nighttime scans from 161 NEXRAD stations during April, May, September, and October during the first 3 hours after local sunset; validation data was selected from April 15–May 15 at 3 hours after sunset from a subset of 16 geographically representative stations. These time periods were selected to coincide with the greatest amounts of bird migration but also included significant bat and insect activity. For example, bat roost emergences followed by widespread foraging were prevalent at several Texas stations, and insect activity was common at many southern stations, with insect-dominated cases occurring on nights that were unfavorable or outside the peak window for bird migration. MistNet was found on this validation set to identify at least 95.9 % of all biological reflectivity with a false discovery rate of 1.3 %.

MistNet has been widely used for aeroecology in the US as part of both WSRLIB and bioRad, and in our experience performs consistently at identifying widespread biological scattering of different taxa. This is compatible with it primarily being trained to identify *precipitation* using a convolutional neural network, which is skilled at recognizing features such as shape, edge structure, and the large vertical distribution of precipitation compared with biological scattering, which is usually concentrated near the ground (e.g., see Figure 7). We are not aware of seasonal or geographical changes in performance, but this has not been systematically evaluated. Based on our own observations, MistNet often misclassifies dense, localized scattering from bird and bat roosts as precipitation, but correctly classifies the same scatterers once they disperse to fill a large area. We expect these errors to have a minor impact for the purposes of this dataset, which quantifies widespread biological scattering within a radius of 50 km, and recommend other approaches for identifying any analyzing roosts [85–88].

### Reflectivity Profiles

We built reflectivity profiles using sampling volumes within a ground distance of 50 km of the radar, with height of up to 3000 m above the radar, and from sweeps at the five elevation angles closest to 0.5*^◦^*, 1.5*^◦^*, 2.5*^◦^*, 3.5*^◦^* and 4.5*^◦^*. Each sweep was first aligned to a fixed 0.5*^◦^* by 250 m polar grid using nearest neighbor interpolation. We then performed clutter, occultation, and precipitation filtering by aligning each sampling volume to the closest entry of each mask and then applying the following filtering rules. Filtering can either set the reflectivity to zero if there is evidence of the absence of biological scatterers (used for precipitation and “NODATA”, described below), or exclude a sampling volume from the set used to calculate average reflectivity if the measurement of biological scattering is judged to be unknown or unreliable (used for clutter and occultation):

- We excluded sampling volumes marked as static clutter or having occultation.
- We excluded sampling volumes with radial velocity between *−*1 m s*^−^*^1^ and 1 m s*^−^*^1^. This is a common practice to mitigate scattering from “dynamic clutter”, i.e., stationary objects that are not captured by static clutter maps [16, 21, 27, 31, 109].
- Reflectivity was set to zero when a sampling volume was identified as precipitation. Although biological scatterers may mix with precipitation, we lack methods to reliably separate biological scatterers in the same sampling volume, and generally assume that biological activity is limited in precipitation, especially when it is intense or widespread.
- Reflectivity was set to zero when the reflectivity factor measurement was “NODATA”. The “NODATA” designation occurs when the received radio waves are below the signal-to-noise threshold, which indicates the absence of sufficient scatterers to produce a detectable signal.

Finally, we computed the average reflectivity in each height bin for the non-excluded sampling volumes. To preserve information about precipitation in addition to biological scattering, we also computed average reflectivity without the precipitation filtering step.

### Cloud Computing

We processed volume scans using Microsoft’s Azure cloud computing service. We created a MATLAB standalone executable to process batches of 100 scans and deployed the executable in a Docker container running Ubuntu Linux and the MATLAB runtime environment. Scans took an average of 11 seconds to process on a single Azure Dv2-series virtual CPU (vCPU). We ran each batch as a separate task using the Azure Batch service. Our compute nodes were Dv2-series virtual machines with two vCPUs and 7GB memory, each running up to two concurrent tasks. Upon completion, the vertical profile for each scan was saved in comma-separated value (CSV) format to an S3 bucket on Amazon Web Services. At 11 seconds per scan, the total compute time for 298 million scans was approximately 104 vCPU years, i.e., the rough equivalent of 104 years on a single CPU.

We took steps to log and recover from several types of processing errors. Errors on some fraction of volume scans are expected. The NEXRAD archive has collected data consistently and uniformly over a long time period, but still has significant variability due to radar hardware and software upgrades, the introduction of new data message types, different compression formats, different radar operating modes, and incomplete or corrupted files. As a result, software may fail or data may be unsuitable for some tasks. When possible, we caught and logged errors that occurred for individual volume scans.

Some Azure compute tasks, corresponding to batches of 100 volume scans, also failed. Some failures were for expected reasons such as the failure or preemption of compute nodes. Others were caused by software crashes, usually in the RSL data ingestion routines and triggered by certain (often corrupted) data files. To overcome errors in compute tasks, we first retried each batch in full to overcome transient issues. We then split each batch of 100 into smaller batches—first 10 scans, and then 1 scan—to attempt to process as many scans as possible while bypassing crashes due to corrupt files.

We successfully constructed profiles for 296.0 million volume scans, which was 99.2 % of the available files. Error messages were logged for 2.27 million scans, of which 84.3 % explicitly indicated a data issue, while 15.7 % indicated an unknown issue. For example, the most common error (47.5 %) was the lack of a data sweep within 1*^◦^* of one of the five requested elevation angles—which was required for precipitation filtering using MistNet and alignment to the clutter and occultation masks—followed by “no velocity data” (16.9 %), “only one sweep available” (11.1 %), and “no reflectivity data” (8.0 %). A total of 278 scans were suspected causes of software crashes based on repeated failures.

### Time-Series Data Products

We constructed several time-series data products by aggregating the vertical profiles. We first vertically integrated each profile to create point measurements for each volume scan. The point measurements include *vertically integrated reflectivity* (VIR) and (vertically integrated) *reflectivity traffic rate* (RTR). VIR (cm^2^ km*^−^*^2^) is the sum of reflectivity (cm^2^ km*^−^*^3^) multiplied by bin height (km) across all height bins. RTR (cm^2^ km*^−^*^1^ h*^−^*^1^) is the sum of reflectivity (cm^2^ km*^−^*^3^) multiplied by speed (km h*^−^*^1^) and bin height (km), and represents the total RCS of scatterers crossing a 1km transect orthogonal to the direction of travel per hour. RTR can be converted to *migration traffic rate*, a standard measure of movement intensity, by supplying information about the RCS of scatterers [89]. We computed VIR and RTR both with and with out precipitation filtering, and computed point measurements of velocity and speed as reflectivity-weighted averages of the corresponding variables over height bins. Reflectivity weighting is used so that each unit of scattering area contributes equally, as opposed to each unit of volume, which may contain different numbers of scatterers. In very rare cases (a few across the entire archive) we found erroneous extremely high velocity measurements due to numerical instabilities in the velocity profiling algorithm, which also led to extremely high RTR values. We excluded these by setting velocity-related variables to missing whenever the speed in an elevation bin exceeded 50 m s*^−^*^1^.

We next assembled point measurements into scan-level time series for each radar station. Each record in the time series corresponds to a single volume scan, so timestamps are irregularly spaced and match the original collection times. For each record we added information about the solar elevation and diel period (night or day) based on the solar position algorithm [115] in the pvlib Python package. Day was defined as the period between local sunrise and sunset, and night was defined as the period between sunset and sunrise on the calendar date. Sunrise and sunset were defined as the times when the solar elevation was *−*0.8333*^◦^*, which, under standard refraction conditions, is the moment when the sun’s upper limb crosses the horizon.

To facilitate downstream analyses that benefit from regular timestamps, such as ones that combine time-series from many stations, we then created 5-minute time series by resampling to a fixed time step of 5 minutes. We resampled data fields using linear interpolation if the time to the nearest observation was less than 1 hour, and left data fields empty otherwise. If the time to the nearest observation was more than 10 minutes but less than one hour, the record was marked as “filled”.

Finally, to facilitate even higher-level analyses, we created daily time series by aggregating the 5-minute time series across three different daily periods: night, day, and the full UTC calendar day. For density variables, we aggregated by integrating across the time period: for example, to integrate reflectivity we summed the reflectivity point measurement (cm^2^ km*^−^*^2^) for each 5-minute interval and multiplied by the interval length (<u>^1^</u> h) to obtain the total reflectivity-hours (cm^2^ km*^−^*^2^ h) for the period. For velocity variables, we aggregated by computing the reflectivity-weighted average across the time period. For details of each variable, see the data record description.

## Data Records

All data records are hosted on Zenodo [116]. Seven data records hold vertical profiles and an eighth holds time series:

- Vertical profiles for 1995–1999 [117]: https://doi.org/10.5281/zenodo.18436894
- Vertical profiles for 2000–2004 [118]: https://doi.org/10.5281/zenodo.18436889
- Vertical profiles for 2005–2009 [119]: https://doi.org/10.5281/zenodo.18436884
- Vertical profiles for 2010–2014 [120]: https://doi.org/10.5281/zenodo.18436881
- Vertical profiles for 2015–2019 [121]: https://doi.org/10.5281/zenodo.18436879
- Vertical profiles for 2020–2024 [122]: https://doi.org/10.5281/zenodo.18436874
- Vertical profiles for 2025–2029 [123]: https://doi.org/10.5281/zenodo.18436969
- Time Series Data from 1995–2025 [124]: https://doi.org/10.5281/zenodo.18433334

In addition, a GitHub repository (http://github.com/darkecology/darkeco-dataset) serves as a central entrypoint to the dataset with links to Zenodo records, download instructions, data documentation, and code examples.

### Vertical Profiles

The profiles for each year are combined in bzip2-compressed archive files (e.g., profiles_1995.tar.bz2) ranging in size from 2.3–9.5 GB and grouped into Zenodo records for 5-year periods starting in 1995. After downloading and extracting the data archive for a given year, there will be one csv file per radar volume scan containing the vertical profile information. Files are grouped into folders by date and station following the same directory structure as the source data (the unidata-nexrad-level2 bucket on AWS). For example, the folder profiles/2017/06/06/KBOX contains profiles from station KBOX on June 6, 2017. This folder includes data files with names such as KBOX20170606_000437.csv, KBOX20170606_000957.csv, and KBOX20170606_001517.csv, each containing one vertical profile. In more detail, the filename format is profiles/YYYY/MM/DD/CCCC/CCCCYYYYMMDD_hhmmss.csv, where YYYY is the four-digit year, MM is the two-digit month, DD is the two-digit day, CCCC is the four-character station identifier (call sign), and hhmmss is the UTC time of the volume scan expressed in hours, minutes, and seconds.

Each vertical profile csv file contains one row per height bin with columns indicating: (1) the height of the bin, (2) the measurements for the bin, including reflectivity and velocity, and (3) metadata for the volume scan, which is repeated in each row. The full schema and description of each field is given in Table 2. Missing values may occur when a height bin contains no scattering volumes for the relevant radar variable or, for velocity-related fields, if numerical issues during velocity profiling prevented estimation.

For readers familiar with the bioRad R package [89], Dark Ecology vertical profiles correspond to vp or vpts objects and can be ingested in bioRad using the read_cajun function for further analysis; see Table 1 for more details.

**Table 1:**
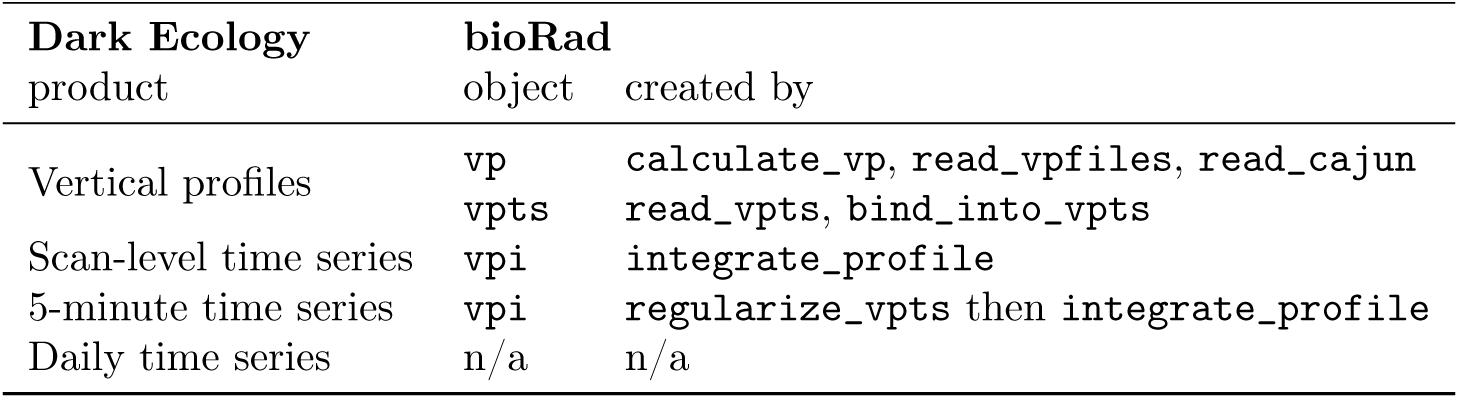
Relationship between Dark Ecology data products and bioRad [89] objects. Relevant bioRad classes are vp (vertical profile), vpts (vertical profile time series), and vpi(integrated profile). The bioRad function read_cajun ingests Dark Ecology vertical profiles as vp objects, which can be bound into vpts objects or vertically integrated to form vpi objects.

**Table 2:** Vertical profile schema. An asterisk indicates a required field.

| Name | Description | Type | Unit |
| --- | --- | --- | --- |
| bin_lower* | Lower limit of height bin. | integer | m |
| height* | Center point of height bin. | integer | m |
| linear_eta | Average linear reflectivity for height bin. Represents the radar cross section in $\text{cm}^2$ per cubic kilometer of atmosphere. | number | $\text{cm}^2 \text{km}^{-3}$ |
| nbins* | Number of scattering volumes in height bin. | integer |  |
| direction | Mean direction of travel. The angle is given as a compass bearing in degrees clockwise from north. | number | degree |
| speed | Mean (ground) speed of travel. | number | $\text{m s}^{-1}$ |
| u | Zonal (east-west) velocity component. | number | $\text{m s}^{-1}$ |
| v | Meridional (north-south) velocity component. | number | $\text{m s}^{-1}$ |
| rmse | Root-mean squared error of VVP fit for scattering volumes in height bin. | number | $\text{m s}^{-1}$ |
| elev1* | Elevation angle of lowest sweep. | number | degree |
| nvolumes_gr35_e1* | Number of scattering volumes with reflectivity factor exceeding 35 dBZ in lowest selected sweep. | integer |  |
| elev2* | Elevation angle of second lowest sweep. | number | degree |
| nvolumes_gr35_e2* | Number of scattering volumes with reflectivity factor exceeding 35 dBZ in second lowest selected sweep. | integer |  |
| vcp* | NEXRAD volume coverage pattern for scan. Examples are 31, 32, 35 (clear air), 12, 212, 112, 215, 121 (precipitation). | integer |  |
| linear_eta_unfiltered | Average reflectivity for all scattering volumes in height bin, including those classified as rain. | number | $\text{cm}^2 \text{km}^{-3}$ |
| percent_rain | Fraction of scattering volumes in height bin classified as precipitation. This fraction is weighted toward regions closer to the radar in accordance to the density of scattering volumes, which are smaller and more densely packed at closer ranges. In the derived data products, this variable is renamed to ‘fraction_rain’. | number |  |

### Time Series

Another Zenodo record holds time series data from 1995–2025 [124]. This record contains separate zip files for the time series at different aggregation levels:

- Scan-Level Time Series (scans.tar.bz2). After unzipping, files are organized into directories by year with one csv file per station-year (e.g. scans/2017/KBOX-2017.csv). Each file has one row per volume scan with fields representing vertically integrated variables from the vertical profiles. The full schema and description of each field is given in Table 3.
- 5-Minute Time Series (5min.tar.bz2). The format is similar to the scan-level time-series, but timestamps are all at 5-minute intervals. After unzipping, there is one csv file per station-year (e.g., 5min/2017/KBOX-2017-5min.csv) with one row per 5-minute interval and fields that are interpolated from the scan-level time series. The full schema and description of each field is given in Table 4.
- Daily Time Series (daily.tar.bz2): There is one csv file per year (e.g., daily/2017-daily.csv). Each file has one row per day and period (day, night, utc_calendar_day) with fields representing aggregated measurements from the 5-minute time series for that period. The full schema and description of each field is given in Table 5.

**Table 3:**
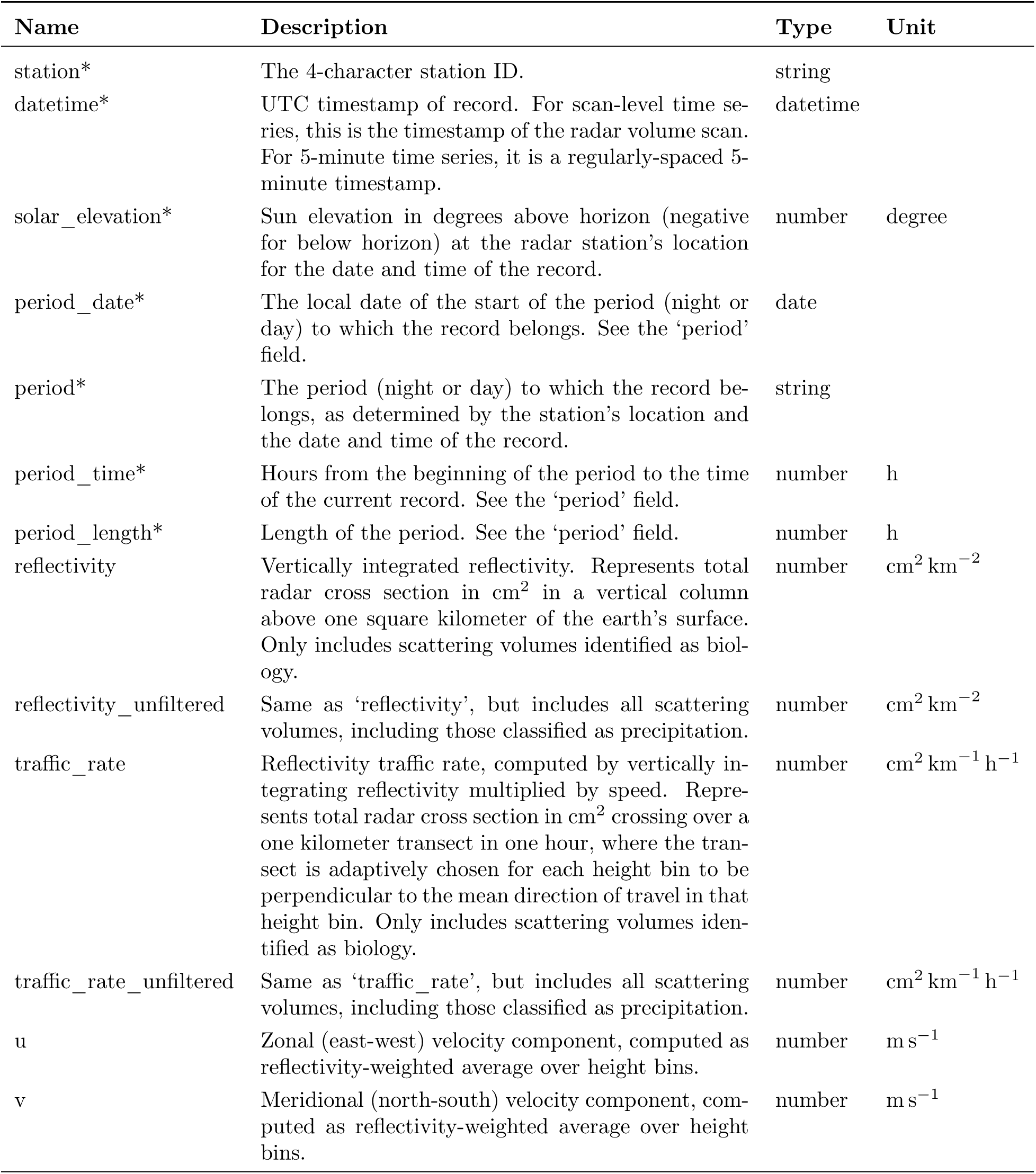
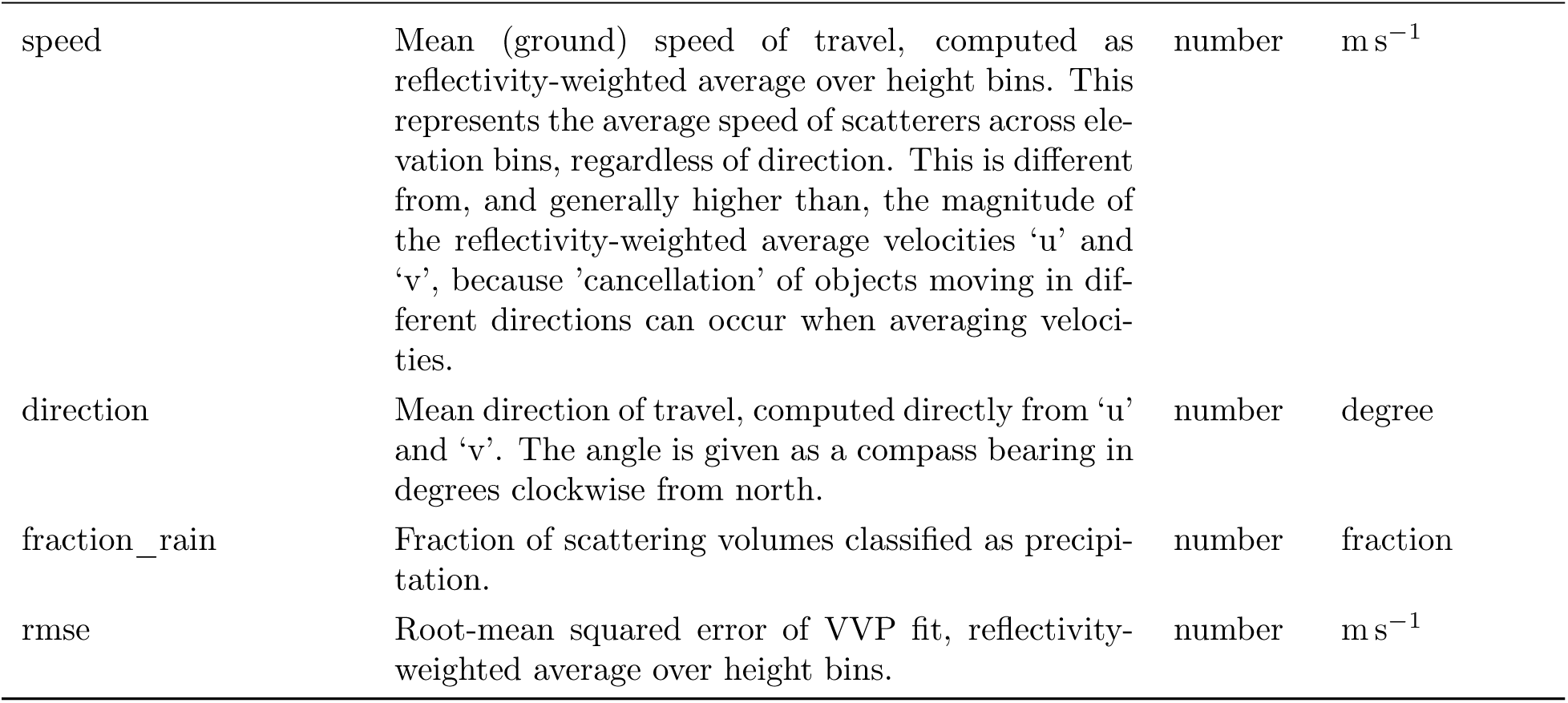
Scan-level time series schema. An asterisk indicates a required field.

**Table 4:**
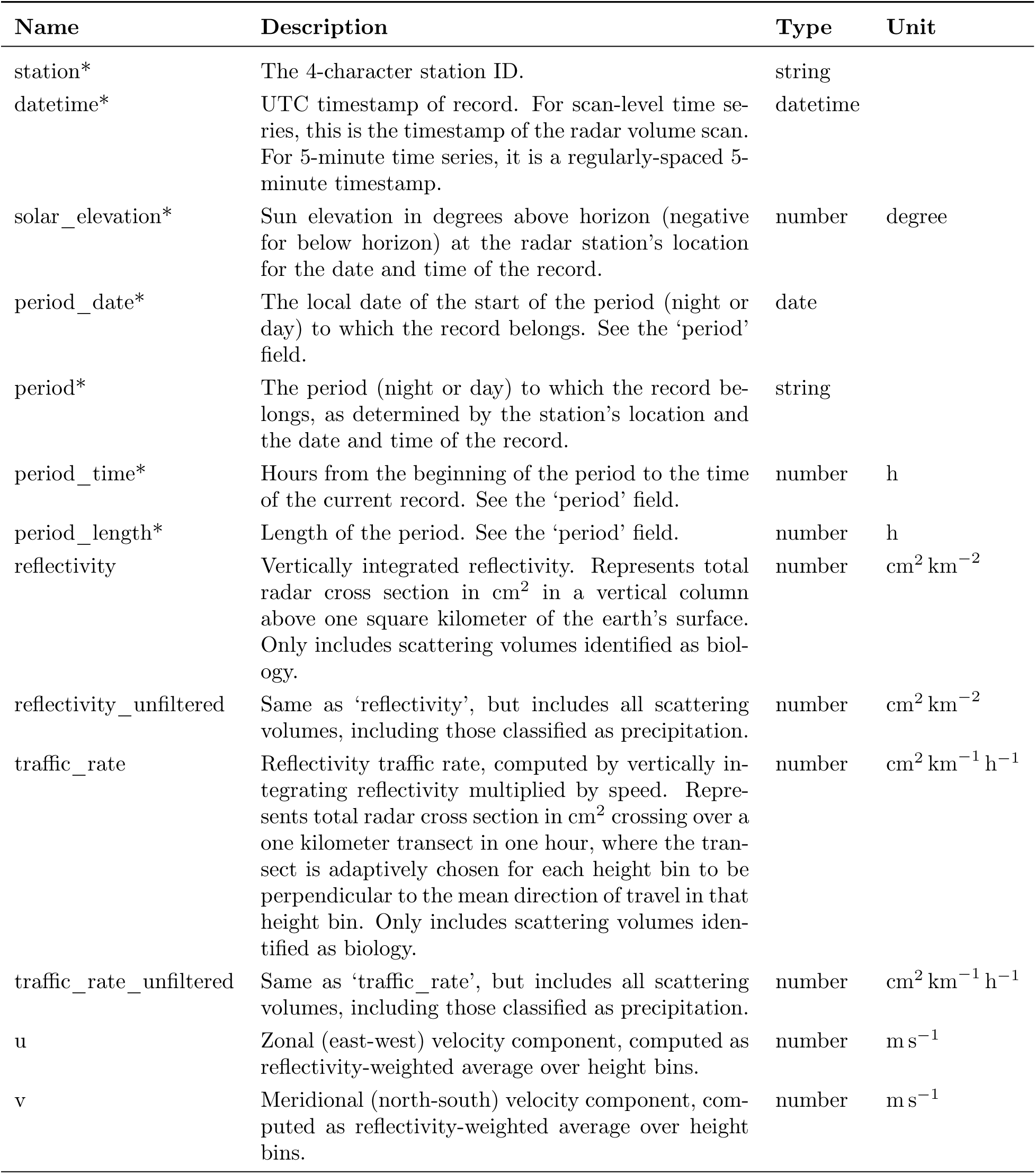
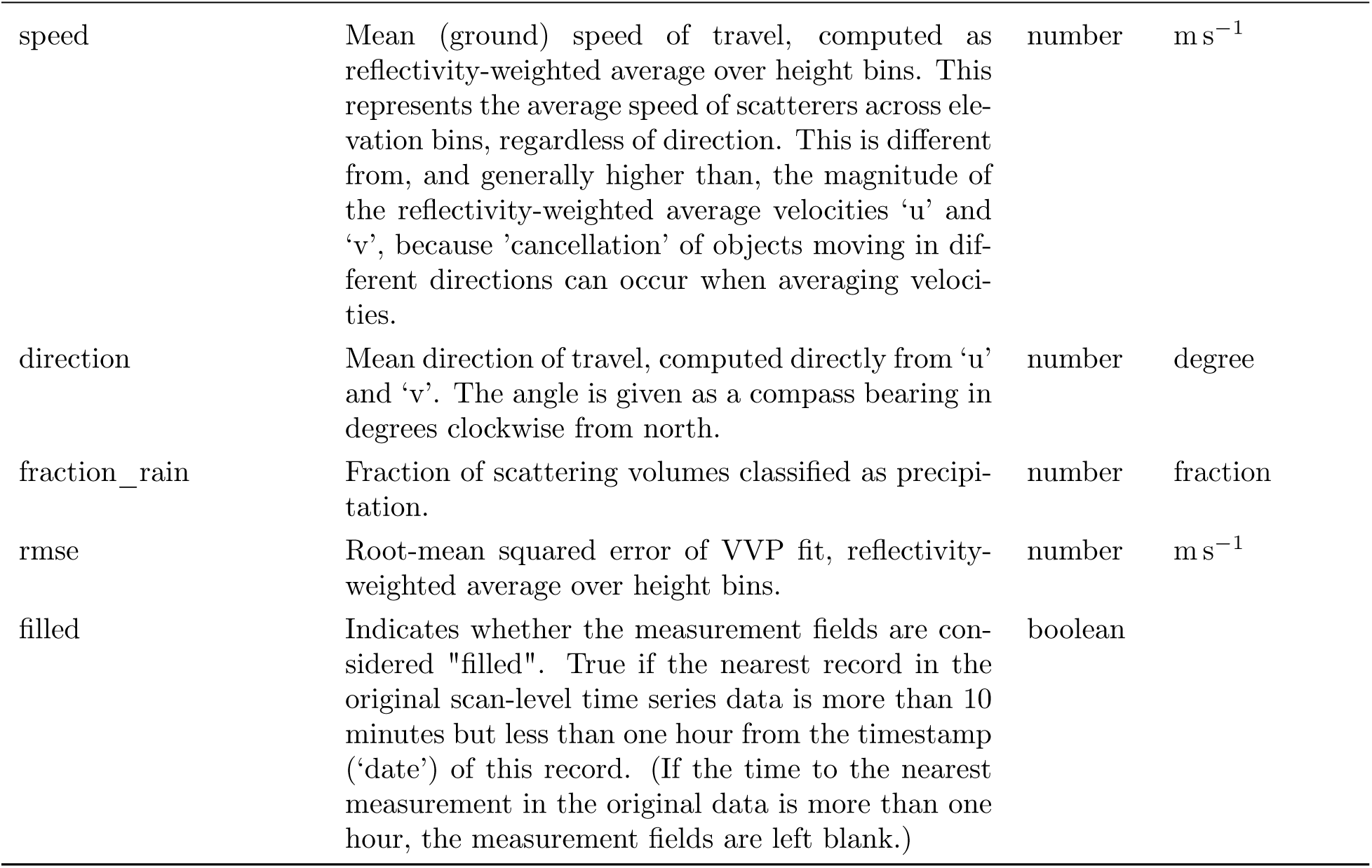
5-minute time series schema. An asterisk indicates a required field.

**Table 5:** Daily time series schema. An asterisk indicates a required field.

| Name | Description | Type | Unit |
| --- | --- | --- | --- |
| station* | The 4-character station ID. | string |  |
| date* | UTC date of the record. | date |  |
| period* | The period of the day ('night', 'day', or 'utc_calendar_day') for which records of the 5-minute time series were aggregated to produce this record. | string |  |
| period_length* | Length of the period. See the 'period' field. | number | h |
| fraction_missing | Fraction of 5-minute records in this period that were 'missing' when aggregating. | number |  |
| fraction_filled | Fraction of 5-minute records in this period that were 'filled' in the 5-minute time series before aggregating. | number |  |
| reflectivity_hours | Temporally and vertically integrated reflectivity. Represents accumulated hours of radar cross section in $\text{cm}^2$ from a vertical column above one square kilometer of the earth's surface. Only includes scattering volumes identified as biology. | number | $\text{cm}^2 \text{ km}^{-2} \text{ h}$ |
| reflectivity_hours_unfiltered | Same as 'reflectivity_hours', but integrates 'reflectivity_unfiltered' over time instead of 'reflectivity', so it includes measurements from all scattering volumes, including those identified as rain. | number | $\text{cm}^2 \text{ km}^{-2} \text{ h}$ |
| traffic | Reflectivity traffic, computed by integrating 'traffic_rate' over time. Represents total radar cross section in $\text{cm}^2$ crossing over a one kilometer transect during the time period. Only includes scattering volumes identified as biology. | number | $\text{cm}^2 \text{ km}^{-1}$ |
| traffic_unfiltered | Same as 'traffic', but integrates 'traffic_rate_unfiltered' over time instead of 'traffic_rate', so it includes measurements from all scattering volumes, including those identified as rain. | number | $\text{cm}^2 \text{ km}^{-1}$ |
| u | Zonal (east-west) velocity component, computed as reflectivity-weighted average over the time period. | number | $\text{m s}^{-1}$ |
| v | Meridional (north-south) velocity component, computed as reflectivity-weighted average over the time period. | number | $\text{m s}^{-1}$ |
| direction | Mean direction of travel, computed as reflectivity-weighted average over the time period. The angle is given as a compass bearing in degrees clockwise from north. | number | degree |
| speed | Mean (ground) speed of travel, computed as reflectivity-weighted average over the time period. | number | $\text{m s}^{-1}$ |
| fraction_rain | Fraction of scattering volumes classified as precipitation, computed as a simple average over the time period. | number |  |

Data schemas for each different type of data file (vertical profiles, scan-level time series, 5-minute time series, and daily time series), written in the Frictionless Table Schema [125], are also included in the associated Zenodo records. A metadata file, nexrad_stations.csv, providing information about each radar station such as its call sign, location, and elevation, is included in each data record for convenience.

Scan-level time series and 5-minute time series correspond to vpi objects in bioRad (Table 1). bioRad does not have a direct analog for daily time series.

### Technical Validation

Core elements of our methods and software have been used in many prior research studies and validated in those contexts [21–23, 29, 31, 32, 34, 39, 41–44, 81, 86]. This section provides additional validation of Dark

Ecology data through (i) a comparison of our reflectivity measurements to those produced by bioRad [89], a widely used tool for aeroecological analysis of NEXRAD data, and (ii) a demonstration that Dark Ecology data can be used to infer coherent patterns of biological activity across space and time that are consistent with prior knowledge.

### bioRad Comparison

#### Profiles

We compared Dark Ecology data to vertical profiles computed by bioRad using the vol2bird algorithm [16]. While there is no established ground truth or gold standard for vertical profiles of biological activity, bioRad is another widely used software for vertical profiles, and comparing these two independent and well tested methods can provide additional confidence in both of their outputs. The relationship between Dark Ecology data products and bioRad objects is summarized in Table 1. We computed vol2bird profiles for the KBGM station in Binghamton, NY from October 1 to 15, 2019 at 15 minute increments. For ease of comparison, we vertically integrated both sets of profiles to scan-level time series (vpi objects) using bioRad’s vertical integration routine.

Figure 4 shows that the reflectivity and traffic rate time series produced by Dark Ecology and vol2bird exhibit near-identical temporal structure, with Dark Ecology yielding slightly lower magnitudes on average. Velocity variables also track each other closely, though with increased variability, particularly during periods of low reflectivity. During periods of high reflectivity— corresponding predominantly to bird migration in this case study—the *u* and *v* velocity components of the two methods agree more closely. Scatter plots of the same variables across this time period are shown in Figure 5. The two methods show extremely strong linear correlation for both reflectivity (0.997) and traffic rate (0.996). Dark Ecology data reports, on average, 90.2 % of the reflectivity and 90.9 % of the traffic rate of vol2bird (linear model fits, *p <* 2 *×* 10*^−^*^16^). These magnitude differences are mostly likely attributed to differences in the spatial ranges used to compute profiles (35 km for vol2bird and 50 km for Dark Ecology), as well as minor differences in filtering steps. The unfiltered *u* and *v* velocities exhibit correlations of 87.8 % and 94.1 %, respectively, and cluster around the 1:1 line. One prominent deviation arises from a behavior where vol2bird estimation defaults to an initial value when the signal-to-noise ratio is low; in this case study, the behavior led to many velocities of exactly 0. When filtering to scans with reflectivity above 100 cm^2^ km*^−^*^2^, the correlations increase to 0.956 and 0.973 for *u* and *v*, respectively, and the clustering around the 1:1 line tightens considerably.

**Figure 4:**
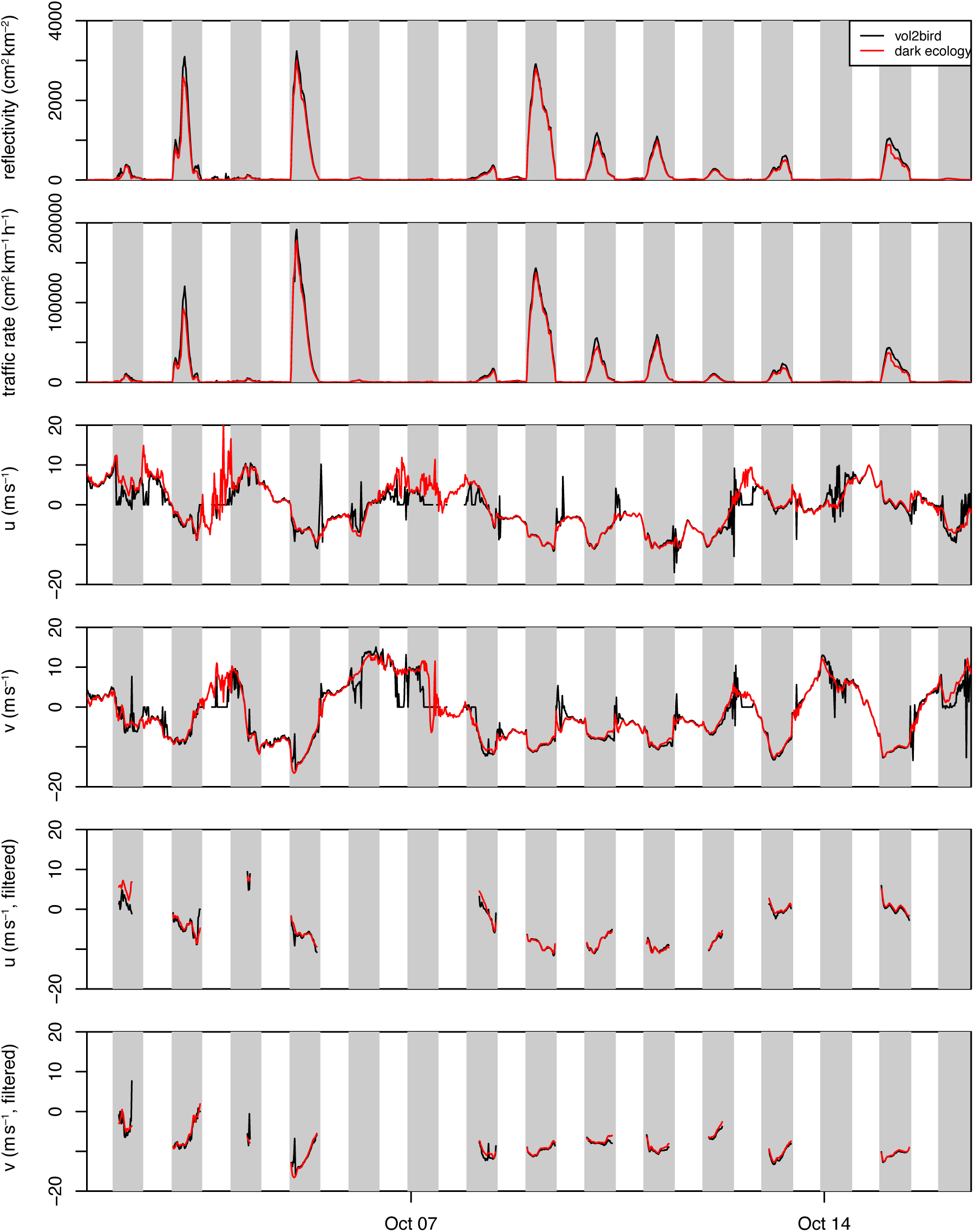
Comparison of scan-level time series obtained by integrating vertical profiles. Variable names follow Table 3. Reflectivity and traffic rate measurements (top two rows) obtained from vol2bird and Dark Ecology are very similar, with vol2bird values slightly higher. Velocity measurements (third and fourth rows) of both methods also follow similar trends but are variable during periods of low reflectivity, when few scatterers are present. Velocity measurements match more closely when filtered to scans where the reflectivity is at least 100 cm^2^ km*^−^*^2^ (bottom two rows), indicative of more biological scatterers being present.

**Figure 5:**
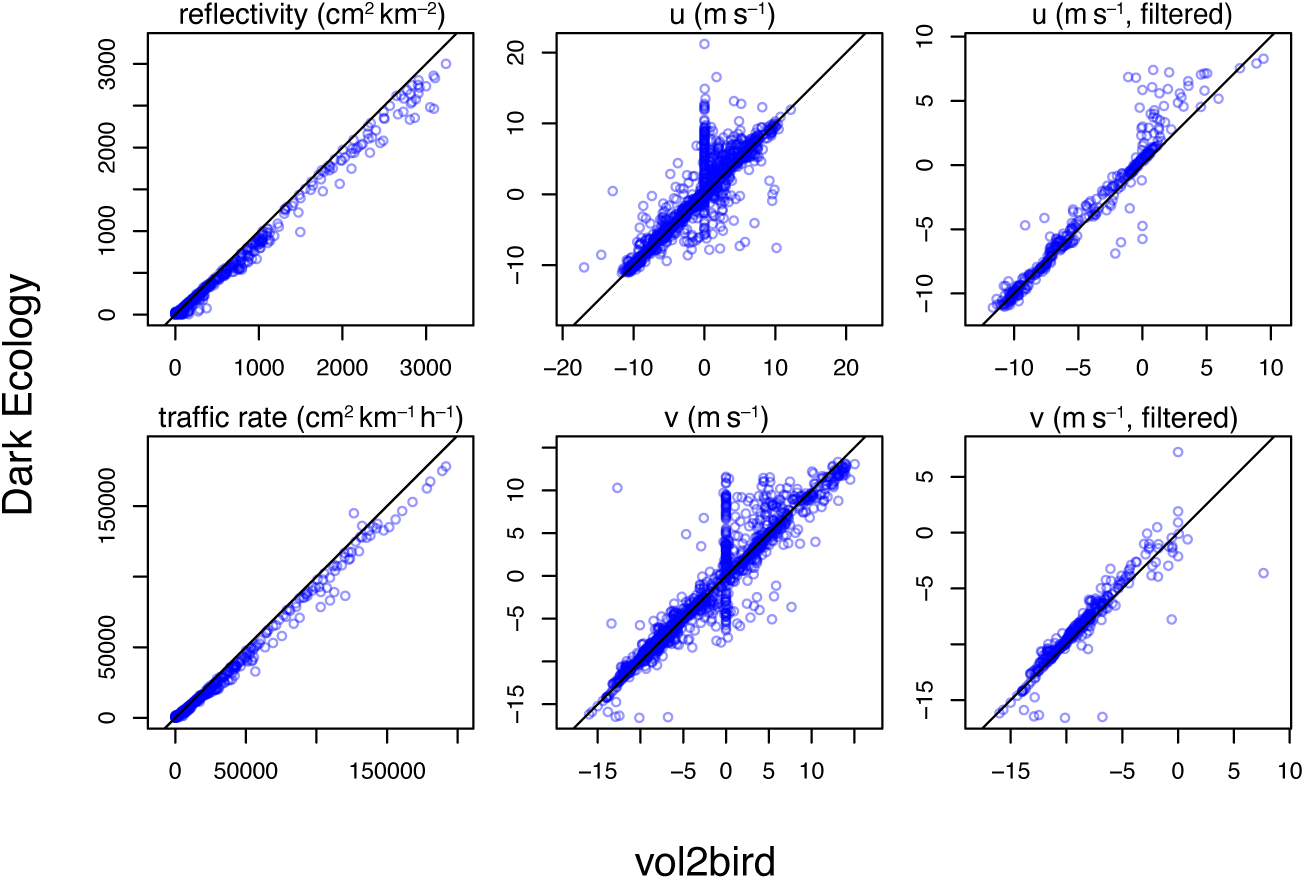
Scatter plots of reflectivity and velocity measurements from Dark Ecology and vol2bird. Variable names follow Table 3. The two data sources are highly correlated for reflectivity and traffic rate (left column), but the Dark Ecology values are typically slightly lower. The *u* and *v* velocity components are also highly correlated (middle column), but with more variability, especially when the reflectivity is low. vol2bird often reports velocities of exactly zero due to a default behavior in the estimation routine. When filtered to scans where the reflectivity is at least 100 cm^2^ km*^−^*^2^ (right column), the velocity variables match more closely.

#### Vertical integration

We also compared vertical integration routines by applying bioRad’s vertical integration method, integrate_profile, to Dark Ecology profiles and comparing the resulting scan-level time series to those produced using our own vertical integration routines. These agreed essentially perfectly, with linear correlation of exactly 1 for all variables and mean average percentage difference (relative to the vol2bird reference value) less than 0.0004 % across all variables.

### Large-Scale Patterns

Analysis of activity patterns in the Dark Ecology data provides strong qualitative validation of the dataset. Figure 6 shows one month of biological reflectivity profiles for the KBGM station in Binghamton, NY, and can be compared with the unfiltered (biology plus weather) profiles in Figure 7. The biological profiles exhibit clear and characteristic signatures of nocturnal bird migration expected at this location and time of year: nightly periods of elevated reflectivity, strongest at lower altitudes, with both intensity and vertical structure varying from night to night. Much lower reflectivity values are observed during the day, with some days showing pronounced diel patterns that are likely due to insects or a mixture of insects and birds. In contrast, the unfiltered profiles display prominent precipitation signatures, appearing as vertical bands of high reflectivity that span the full altitude range. These precipitation artifacts are effectively removed from the biological profiles by the MistNet algorithm, while preserving the underlying biological signal. Visualizations of this kind—high temporal resolution profiles spanning extended time periods—are made easy by the Dark Ecology dataset and provide a compelling and interpretable view of biological phenomena in radar data.

**Figure 6:**
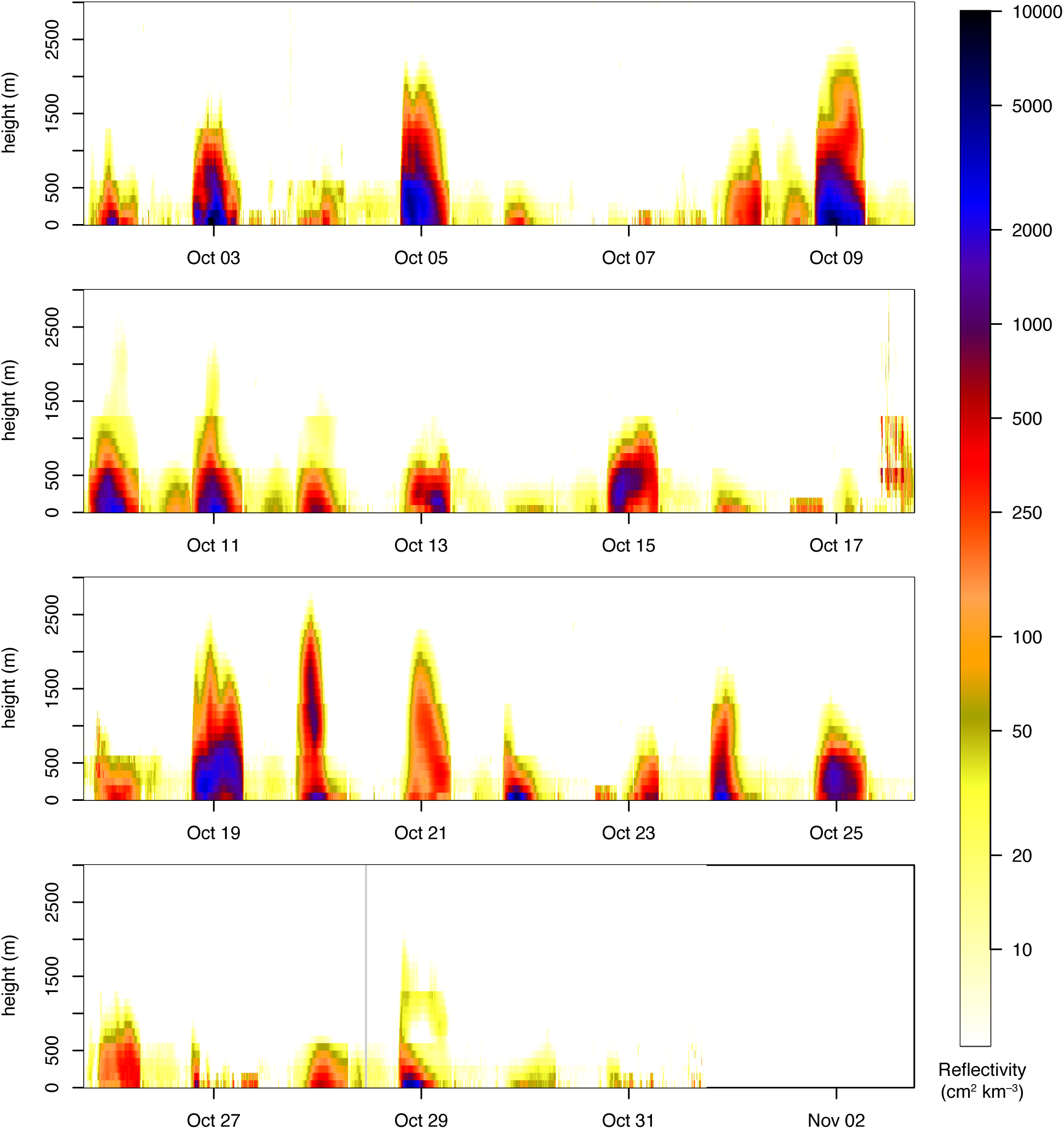
Biological reflectivity profiles for the month of October 2019 at the KBGM radar station in Binghamton, NY. The nightly pattern of bird migration is evident by the periods of increased reflectivity at night with higher intensity at lower elevations and night-to-night variation in the overall intensity and height distribution of migratory activity. Activity peaks (e.g., October 3, 5, and 9) can be matched to those in the the scan-level time series in Figure 4 (top row). Much lower reflectivity values are observed during the day with pronounced diel patterns observable on some days; these may be due to insects or a mixture of insects and birds.

**Figure 7:**
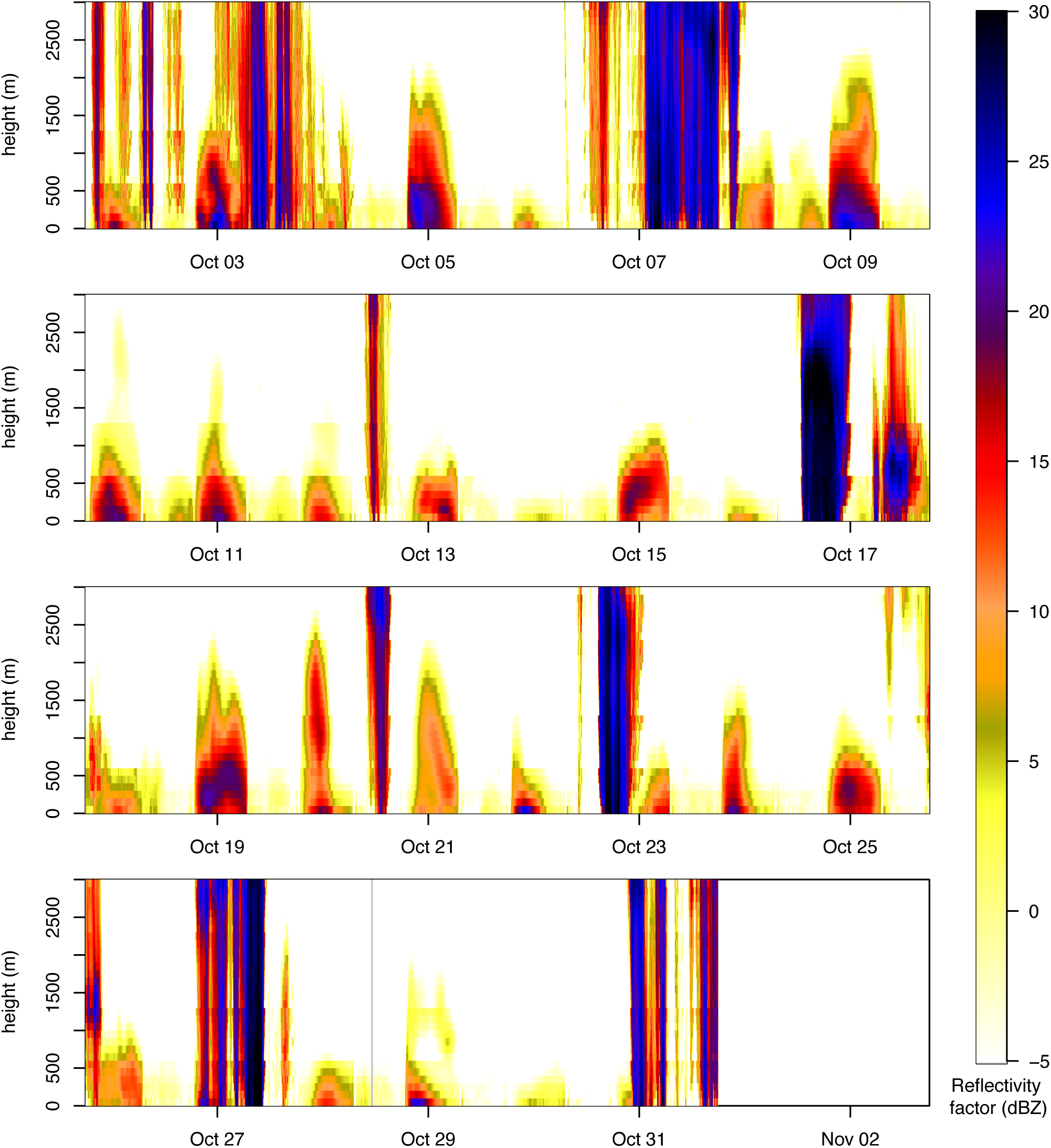
Unfiltered reflectivity profiles, including biology and precipitation, for the month of October 2019 at the KBGM radar station in Binghamton, NY. Intermittent periods of rain appear as vertical stripes of high reflectivity, which, unlike scattering from bird migration, spans across all heights. The visual comparison to Figure 6 shows that rain is effectively removed from the biological profiles by the MistNet algorithm. The color scale is identical to Figure 6, but the values are shown as reflectivity factor, which is more typical for precipitation.

NEXRAD data has seen considerable use for quantifying nocturnal bird migration at broad scales by examining biological reflectivity at night in the spring and fall, when birds are expected to dominate the airspace [27, 31, 37–39, 42]. Figure 8 demonstrates that Dark Ecology data can reveal broad spatial and temporal patterns of bird migration that are consistent with prior knowledge with minimal analyst effort. We focus on the spring season, during which insects and bats have been estimated to contribute minimally to nocturnal biological reflectivity [38]. The spatial distribution of cumulative spring migration traffic (Figure 8a) shows continent-scale patterns that align with known variation in migratory intensity such as a high-intensity corridor in the central US in spring [37]. Temporal summaries of migration phenology show a clear trend of later migration at higher latitudes (Figure 8b) and significant variability across years at a single station (Figure 8c), which is expected due to known correlations between phenology and yearly temperature [39]. At finer temporal scales, daily migration traffic measurements from two nearby stations (Figure 8d) reveal that a large fraction of migration occurs on a small number of nights [42] and that nightly migration intensity is highly correlated at nearby stations, likely due to similar synoptic weather patterns, which are known to be strongly associated with migration [31]. Together, these examples show that Dark Ecology enables exploration of migration dynamics—from continental patterns to individual nights—yielding results consistent with our emerging understanding of avian migration at the largest scales.

**Figure 8:**
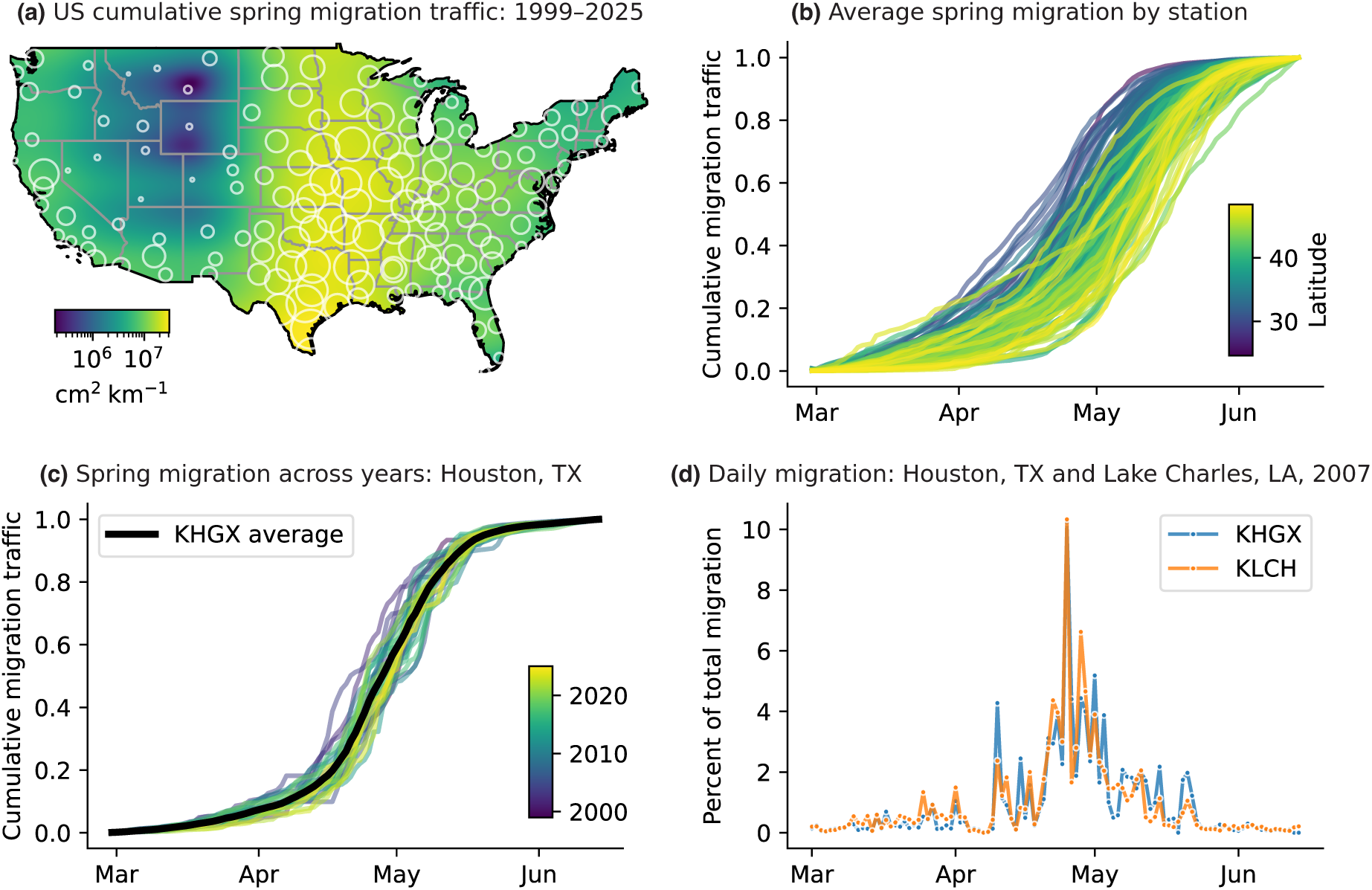
Spatial and temporal structure of nocturnal bird migration in the spring season inferred from Dark Ecology data: (a) Average cumulative spring migration traffic (1999–2025) for each radar station (white circles) with a spatially smoothed estimate shown as a continuous surface. (b) Average cumulative spring migration phenology by station, expressed as the fraction of total seasonal migration completed by date and averaged across years, with curves colored by station latitude. (c) Interannual variability in spring migration phenology at station KHGX (Houston, TX), showing cumulative migration for individual years and the multi-year average (black line). (d) Daily migration traffic in spring 2007 for two nearby stations (KHGX, Houston, TX and KLCH, Lake Charles, LA), expressed as a percentage of each station’s seasonal total.

## Usage Notes

### Software Resources

Several resources are available for working with Dark Ecology data. The main dataset repository [126] includes a folder structure for working with the data as well as download instructions, metadata, links to documentation, and starter code for using the data. The dataset preparation repository [127] includes additional code examples, including scripts to produce figures and analyses from the Technical Validation section of this paper.

The bioRad R package [89] provides tools for biological analysis of radar data and can read Dark Ecology vertical profiles using the function read_cajun. This provides access to bioRad plotting routines and is the easiest way to visualize profile data as in Figures 6 and 7.

### Sources of uncertainty

Users must be aware of the inevitability of errors including the possibility of significant outliers in Dark Ecology data. The NEXRAD network has operated for more than 30 years to collect hundreds of millions of volume scans and many more than 10^15^ individual measurements from across the globe. A sensor network operating at this scale will encounter a huge range of conditions, both in terms of scatterers present in the atmosphere as well as hardware, software, and operational issues. Our data preparation pipeline includes a number of conservative steps to mitigate the effects of clutter and other undesired signals, trim extreme values, and avoid processing data that is incomplete or corrupted. However, it is inevitable that these safeguards are imperfect: rain or clutter may pass our filters or rare radar issues may go undetected. In the following, we briefly discuss several known sources of uncertainty: (1) radar miscalibration or operational issues, (2) loss of vertical resolution due to beam broadening, (3) imperfect filtering of precipitation and other non-biological signals, (4) errors in velocity estimation, (5) changes over time due to radar system upgrades. This list is not exhaustive.

Radar calibration involves determining system parameters such as noise and antenna gain to correctly measure reflectivity. Miscalibration manifests as an additive bias in decibels of reflectivity, or, equivalently, reflectivity that differs from the truth by a constant multiplicative factor. Calibration depends on parameters that may vary over time at different rates [128]. The specified calibration accuracy for NEXRAD is *±*1 dB (i.e., within a factor of 1.26), maintained frequently through online and offline checks [128]. However, comparisons among NEXRAD radars, spaceborne radars, and rain gauge data, mostly made in the early years of NEXRAD (before 2000), have estimated calibration differences for some sites that may be larger and drift or jump suddenly over time [129]. For example, Anagnostou et al. (2001) estimated some radar-to-radar differences to be as high as 2.4 dB (a factor of 1.74) and changes over time at a single site of about 3 dB (a factor of 2.00) [130]. In general, radar calibration has been judged to be an important source of uncertainty for precipitation quantification [129], and we expect that this also holds true for quantifying biological reflectivity in an absolute sense.

The most significant operational issue we are aware of is a very small number of days where radar issues have led to extremely high reflectivity measurements at individual stations. Figure 9 shows an example issue leading to extreme outliers in biological reflectivity measurements at station KDFX on October 3, 2015. We have identified three other similar cases in the dataset: at station KBBX on January 11–12, 2012; at station KBBX on September 12–13, 2012; and at station KPBZ on July 11–12, 2015, and station KDFX on October 3, 2015. This list should not be considered exhaustive. Several other cases of high daily reflectivity measurements were investigated and we could not determine whether they were due to radar issues or actual biological phenomena; more subtle radar operational or calibration issues may be difficult to detect. While users should be aware of these issues, given the very large number of stations and days in the dataset, such extreme outliers should be considered very rare.

**Figure 9:**
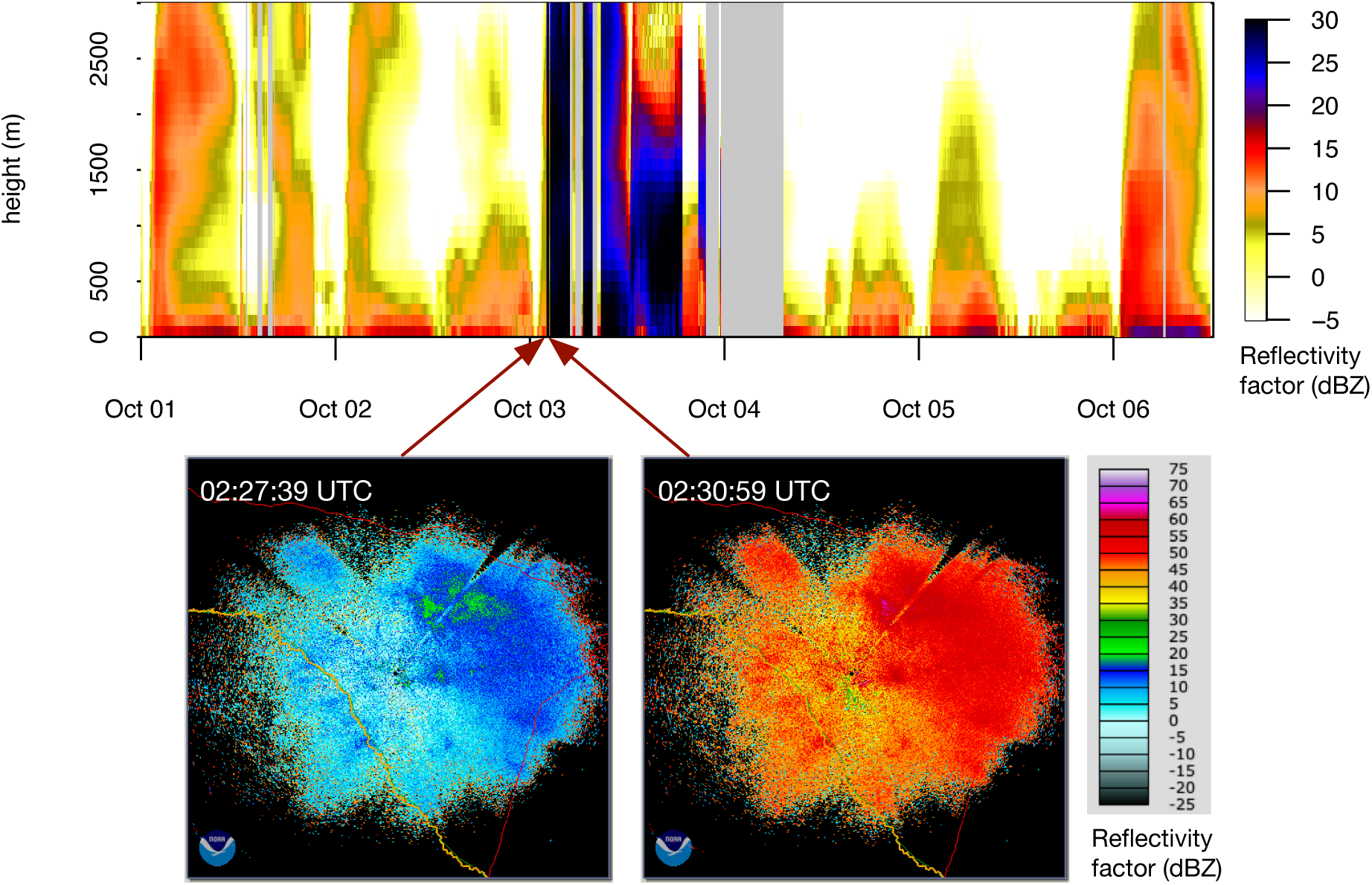
Example of radar operational issue leading to unusually high measurements of biological reflectivity at KDFX (Laughlin Air Force Base, Texas) on October 3, 2015. Vertical profile data (top) shows periods of very high reflectivity with abrupt starts and ends and associated periods of missing data (gray) suggesting the radar was taken offline. The spatial pattern of reflectivity in the 0.5*^◦^* sweep is nearly identical immediately before and after the increase (bottom), but values increased by roughly 30 dBZ, indicating a radar operational issue and not an actual large increase in scatterers (images from NOAA Weather and Climate Toolkit [131]).

Radar geometry and the fact that the beam rises and widens with distance (Fig. 3) affects the interpretation of vertical profiles. By including sampling volumes up to 50 km from the radar, we obtain good coverage at higher altitudes but also experience beam widening that limits the vertical resolution and causes some smoothing relative to the true vertical structure of biological activity. The effects are most pronounced in higher elevation bins, which include sampling volumes at greater distances from the radar. For our spatial domain we estimate the effect to be similar to smoothing the true vertical reflectivity profile with a Gaussian kernel with standard deviation 50–250 m, with less smoothing at low elevations and more at high elevations.

Reflectivity filtering steps are subject to errors, such as misclassifying precipitation as biological scatterers or vice versa. This may occur with any filtering step, but we are most aware of some cases of precipitation contamination: we recommend comparing filtered and unfiltered data (cf. Figures 6 and 7) as a first step to assess this. Precipitation events tend to appear as high reflectivities across a large altitude range (see Figure 7). Unusual signals in the filtered data during precipitation events should be treated cautiously and not over-interpreted. Cases have also been identified where MistNet screens out too much biology, but these cases are rare and seem associated with higher spatial structure and patterning in the biological signal (e.g. KMQT radar, 2020-09-17 00:50:36). Some of our clutter filtering steps are conservative; because these sampling volumes are excluded and not set to zero, the result is a smaller number of sampling volumes used to estimate the average reflectivity in each height bin, as opposed to a systematic reduction in reflectivity values.

Velocity estimation, which affects traffic rate measurements, is also subject to errors, but we do not consider them a major factor for most biological analyses. Elevation bins routinely contain on the order of 10 000 or more sampling volumes; when populated with scatterers, this allows estimating the mean velocity with low variance. On the other hand, velocity measurements associated with low reflectivity values may be based on few scatterers and should not be over-interpreted. Our vertical integration and time series aggregation routines weight by reflectivity, which appropriately limits the impact of noisier measurements in downstream analyses. Velocity estimation also depends on potential errors due to aliasing. For biological scatterers, aliasing is relatively uncommon in NEXRAD data and can be addressed effectively when it occurs. Aliasing occurs most frequently in volume coverage pattern (VCP) 31, a clear air scanning mode that is used about 3 % of the time. This mode has Nyquist velocity of about 12 m s*^−^*^1^, while other VCPs have Nyquist velocity 26 m s*^−^*^1^ or higher, exceeding typical speeds of biological scatterers [132–134]. Even in VCP 31, biological aliasing tends not to be extreme, with most sampling volumes folded at most once, and we have observed very good dealiasing performance [81]. Velocity ambiguity due to aliasing may be more of an issue in C-band radars operating at lower Nyquist velocities [135].

Several upgrades to the NEXRAD network have occurred over the years, which should be considered carefully when conducting historical analyses with the Dark Ecology dataset. In 2007–2008, the “super-resolution” upgrade increased the resolution of sampling volumes from 1*^◦^* to 0.25*^◦^* in azimuth and from 1 km to 250 m in range. In 2012–2013, the software and hardware were upgraded to collect additional dual polarization data products. Both upgrades were designed to maintain consistency in measurements; however, systematic changes in reflectivity measurements of biological scatterers cannot be ruled out, and an earlier analysis found some evidence of a systematic effect from the dual-polarization upgrade [38]. Many other improvements and changes have been made over the years to software, hardware components, algorithms, and operation modes [136–138]. A significant change was the open radar data acquisition (Open RDA) update, completed in 2006 [137], which, among other changes, introduced a new Doppler spectrum-based clutter filtering algorithm [139, 140]. Older NEXRAD data may have significantly more clutter, and changes in the clutter filtering approach may have systematically affected reflectivity measurements [140] (e.g., by reducing bias).

Beyond uncertainties in the Dark Ecology data themselves, we anticipate significant sources of uncertainty associated with downstream uses relating to spatial interpolation, conversions to numbers or biomass, and counting through space and time. For example, one might ask what was the total number or mass of birds (or bats or insects) aloft in a given region. Answering this requires: (1) interpolating or extrapolating from the source measurements, which summarize biological scattering within 50 km of each radar site, to the area of interest, and (2) converting from reflectivity units to numbers or mass units. Both steps involve significant uncertainty because the data do not resolve details of the spatial distribution between radar sites or the identities of scatterers and their RCS or mass. Additional uncertainties may arise with cumulative counting, such as counting the total number of birds that fly within a region in one night, which depends on behavioral characteristics such as speed and duration of flight, which determine how long each scatterer remains in a region of interest. For these types of applications, we suspect that the uncertainty of interpolation, RCS estimation, and other factors will be at least as great as that of the underlying measurements.

### Recommendations

As with all analyses, data users must take their own precautions to ensure validity of their findings. These may include investigating and removing outliers, conducting sensitivity analyses, or employing robust statistical methods. With such measures, we have found that NEXRAD and Dark Ecology data are suitable for a range of biological analyses. The best tool for safeguarding scientific conclusions is familiarity with the data at a detailed level. Although we have provided higher-level data products such as time series for convenience, users are strongly encouraged to become familiar with the vertical profile data as well as the source data (NEXRAD Level II) and the methods for processing it. Visualizing profile data as in Figures 6 and 7 is an excellent way to root out potential data issues and also to reveal truths about patterns of biological scatterers, many of which are not fully explored. Level II radar data can be visualized with a variety of tools; NOAA’s Weather and Climate Toolkit [131] is excellent for interactive data exploration.

Because Dark Ecology data do not distinguish different types of biological scatterers, users must base conclusions about taxa on other information. Most works rely primarily on activity patterns, with the fairly simple interpretation that nocturnal scattering during avian migration is strongly bird dominated [21, 38, 39] while diurnal scattering is typically insect dominated [55]. Bats are common in nighttime data from the southern US from spring through fall [56–58] but their overall populations are estimated to be much smaller than migrating birds in the same areas [38]. Various criteria have been applied to safeguard bird analyses from insect contamination or vice versa—based on airspeed [13, 21, 31], residual error of VVP fit [16, 39], or differential reflectivity [55]—but with unclear impacts. Van Doren and Horton found that airspeed-based insect filtering did not significantly alter continental-scale bird migration forecasts [31]. Precautions are most important when different taxa likely co-occur in large numbers, such as co-occurrence of birds and insects common during daytime. In these cases, we recommend pairing profile data with information on winds aloft to analyze patterns in self-propelled airspeed, which can be informative in assessing relative proportions of fast-flying birds and slower flying insects [141, 142].

Historical analyses of trends in abundance or biomass over time must consider the potential impacts of the system changes and upgrades mentioned above. The most extensive prior analysis is by Rosenberg et al. [38], who excluded data prior to 2007 due to the unknown quality and changes in clutter filtering, and included in their statistical models the possibility of systematic changes due to super-resolution and dual-polarization upgrades—they found little evidence of an effect from the super-resolution upgrade and weak evidence of an effect from the dual-polarization upgrade. We advise users of the Dark Ecology dataset to use similar caution. Users must also take care to deal appropriately with missing data, which is significantly higher before about 2004.

Analyses of phenology are likely more robust because they typically depend only on a normalized measure of the timing of biological activity within each year and not the total amount of scattering. For example, a common approach is to extract dates from normalized curves of cumulative activity such as those in Figure 8 [39].

## Data availability

All data is archived in Zenodo [116]:

Vertical profiles for 1995–1999: https://doi.org/10.5281/zenodo.18436894
Vertical profiles for 2000–2004: https://doi.org/10.5281/zenodo.18436889
Vertical profiles for 2005–2009: https://doi.org/10.5281/zenodo.18436884
Vertical profiles for 2010–2014: https://doi.org/10.5281/zenodo.18436881
Vertical profiles for 2015–2019: https://doi.org/10.5281/zenodo.18436879
Vertical profiles for 2020–2024: https://doi.org/10.5281/zenodo.18436874
Vertical profiles for 2025–2029: https://doi.org/10.5281/zenodo.18436969
Time Series Data from 1995–2025: https://doi.org/10.5281/zenodo.18433334

Users are encouraged to access the data through https://github.com/darkecology/darkeco-dataset, which provides the recommended directory structure, download instructions, documentation, and code examples.

## Code availability

The WSRLIB MATLAB toolbox [91] used to read and process NEXRAD Level 2 data to construct vertical profiles is available at https://github.com/darkecology/wsrlib and archived at https://doi.org/10.5281/zenodo.18499707. Code to process vertical profile data and produce time series and to produce figures and analyses in the Technical Validation section is available at https://github.com/darkecology/darkeco-dataset-prep and archived at https://doi.org/10.5281/zenodo.18499923.

## Supplementary materials

Table S1: NEXRAD stations included in the analysis, including, location, elevation, number of years active, and number of scans. Of the 161 stations included in this study, 158 are currently operational and three (KJAN, KLIX, and RODN) have been retired or replaced.

## Acknowledgments

We are grateful to Steve Kelling for helping secure initial funding for the Dark Ecology project and managing the early Dark Ecology research efforts at Cornell.

## Author contributions statement

DS, AD, KH, AF, FAL, and SM conceived the study. DS and SM managed the study. DS, KW, GB, PB, TL, and KH developed software to process data. TL and SM developed computer vision methods. DS, ID, PB, and KW developed cloud workflows. DS and KW processed the data. AD, KH, CN, BMV, AF, and FAL contributed expertise on biological interpretation of radar data. PD consulted on the data organization, access policies, schemas, and metadata. DS wrote the manuscript. KH contributed to figure preparation. All authors reviewed and edited the manuscript.

